# Incorporating Radiopacity into Implantable Polymeric Biomedical Devices for Clinical Radiological Monitoring

**DOI:** 10.1101/2023.01.06.523025

**Authors:** Kendell M Pawelec, Ethan Tu, Shatadru Chakravarty, Jeremy ML Hix, Lane Buchanan, Legend Kenney, Foster Buchanan, Nandini Chatterjee, Subhashri Das, Adam Alessio, Erik M Shapiro

## Abstract

Longitudinal radiological monitoring of biomedical devices is increasingly important, driven by risk of device failure following implantation. Polymeric devices are poorly visualized with clinical imaging, hampering efforts to use diagnostic imaging to predict failure and enable intervention. Introducing nanoparticle contrast agents into polymers is a potential method for creating radiopaque materials that can be monitored via computed tomography. However, properties of composites may be altered with nanoparticle addition, jeopardizing device functionality. This, we investigated material and biomechanical response of model nanoparticle-doped biomedical devices (phantoms), created from 0-40wt% TaO_x_ nanoparticles in polycaprolactone, poly(lactide-co-glycolide) 85:15 and 50:50, representing non-, slow and fast degrading systems, respectively. Phantoms degraded over 20 weeks in vitro, in simulated physiological environments: healthy tissue (pH 7.4), inflammation (pH 6.5), and lysosomal conditions (pH 5.5), while radiopacity, structural stability, mechanical strength and mass loss were monitored. The polymer matrix determined overall degradation kinetics, which increased with lower pH and higher TaO_x_ content. Importantly, all radiopaque phantoms could be monitored for a full 20-weeks. Phantoms implanted in vivo and serially imaged, demonstrated similar results. An optimal range of 5-20wt% TaO_x_ nanoparticles balanced radiopacity requirements with implant properties, facilitating next-generation biomedical devices.

## 1. Introduction

The use of implanted biomedical devices has grown rapidly in recent decades. In particular, the application of polymers as medical devices has greatly increased, as a result of their excellent biocompatibility and materials properties, including favorable mechanics and tunable biodegradation profiles. Despite their ubiquitous use in the clinic, implants made from polymers fail for a number of reasons, such as wear, tearing, migration and infection.^[1]^ With the inherent risk of failure following implantation of biomedical devices and to prevent situations that could irreparably impact patient health, there exists an increased need for a clinical methodology for in situ monitoring of device status following implantation.^[1]^ However, for the majority of polymer implants, there is no robust endogenous contrast mechanism for clinical diagnostic imaging, and hence no mechanism for radiologists to diagnose problems prior to catastrophic failure.

Incorporating radiological monitoring of biomedical devices as a standard part of surgical follow-up could be a major step forward in the prevention of emergency interventions.

To impart contrast-generating properties into polymer devices requires modification of or incorporation of polymers with contrast agent specific to a clinical imaging modality.^[2,3]^ Of these clinical modalities, computed tomography (CT) is a widespread technique that utilizes x-rays to form high-resolution 3D maps of tissue. A drawback to CT is that it cannot easily distinguish between different soft tissues with the same sensitivity as other imaging techniques such as magnetic resonance imaging (MRI), and CT exposes patients to small amounts of radiation, which should be limited over a patient’s lifetime.^[4]^ However, for applications that require diagnosing implant damage, without concurrent interrogation of soft tissue status, the high clinical throughput, low-cost and favorable signal-to-noise ratio from surrounding tissue make CT beneficial for in situ monitoring.^[3]^

To utilize CT for monitoring polymeric biomedical devices, radiopacity must be introduced. Chemical modification of the polymer backbone has been proposed, but aside from changes to materials properties, there is the possibility of releasing cytotoxic elements during degradation.^[5,6]^ Introduction of radiopaque nanoparticles are an extremely versatile alternative.^[7,8]^ Tantalum oxide (TaO_x_) nanoparticles, in particular, are biocompatible in vivo with superior CT contrast over traditional iodinated compounds ^[9,10]^, and can further be incorporated into polymeric matrices for use as biomaterials.^[11,12]^ However, the incorporation of nanoparticles into polymeric devices requires study on the impact of materials properties that could alter device function, particularly as the percentage of nanoparticles increases. Specifically, imaging functionality should not interfere with mechanical stability of devices while allowing tracking of features at a scale necessary to diagnose clinical outcomes.

In creating composite materials for biomedical devices, it is not only the incorporation of nanoparticles that can influence materials properties, but also the physiological environment of implantation. A particularly acidic environment, for example one experiencing chronic inflammation or within a tumor ^[13]^, may potentially accelerate the release of nanoparticles and induce local cytotoxicity. For this reason, the degradation of biomaterials after implantation has remained difficult to predict, even in systems where degradation mechanisms have been studied extensively.^[14,15]^ Traditional degradation studies are conducted to simulate the physiological environment (saline, pH 7.4, 37°C), but these have not proved predictive for the complex in vivo environment.^[16]^ As in situ imaging relies on the sustained presence of nanoparticle contrast agent regardless of conditions, polymer degradation and potential nanoparticle release must be characterized in a comprehensive range of physiological environments, from lysosomal conditions (pH 5.5) to healthy tissue (pH 7.4).

The current study demonstrates the use of CT for in situ monitoring of model implantable biomedical devices (phantoms), over 20 weeks, using two of the most common polymers for manufacturing biomedical devices: polycaprolactone (PCL) ^[17]^ and poly(lactide-co-glycolide) (PLGA). ^[18]^ Hydrophobic TaO_x_ nanoparticles were incorporated (0-40wt%) into phantoms, as it has high radiopacity for its relative size and has been formulated to exhibit homogeneous mixing into hydrophobic polymer systems.^[3,11]^ Phantoms were made using PCL, PLGA 50:50 and PLGA 85:15, mimicking porous isotropic scaffolds used for generic tissue engineering applications, covering the full range of predicted degradation behaviors from virtually non-degrading (years), slow degradation (months) and fast degradation (weeks).^[15,17]^ Degradation in phantoms was characterized by tracking structural integrity, mechanics, mass loss and radiopacity. Significant changes in degradation profiles were noted with buffer pH and TaO_x_ incorporation, defining a practical limit of radiopaque contrast agent incorporation to guide future biomedical device design.

## 2. Results & Discussion

As the goal is to incorporate imaging functionality into implanted devices, radiopaque phantoms were manufactured to mimic state of the art tissue engineering implants. Specifically, the phantoms incorporated porosity at two different scales: macro-pores (200-400 μm) and micro-porosity (< 100μm), **Figure 1**(a-c). The interconnected macro-porous structure is designed to facilitate processes like cellular infiltration and the micro-porosity contained within the pore walls can allow the diffusion of nutrients or chemokines within the structure. Both scales of porosity were introduced to FDA-approved biocompatible polymers using salt leaching, a process where polymer solutions are cast around water soluble place holders that are then removed to become pores, which is known to create interconnected and permeable implants.^[19]^

**Figure 1.**
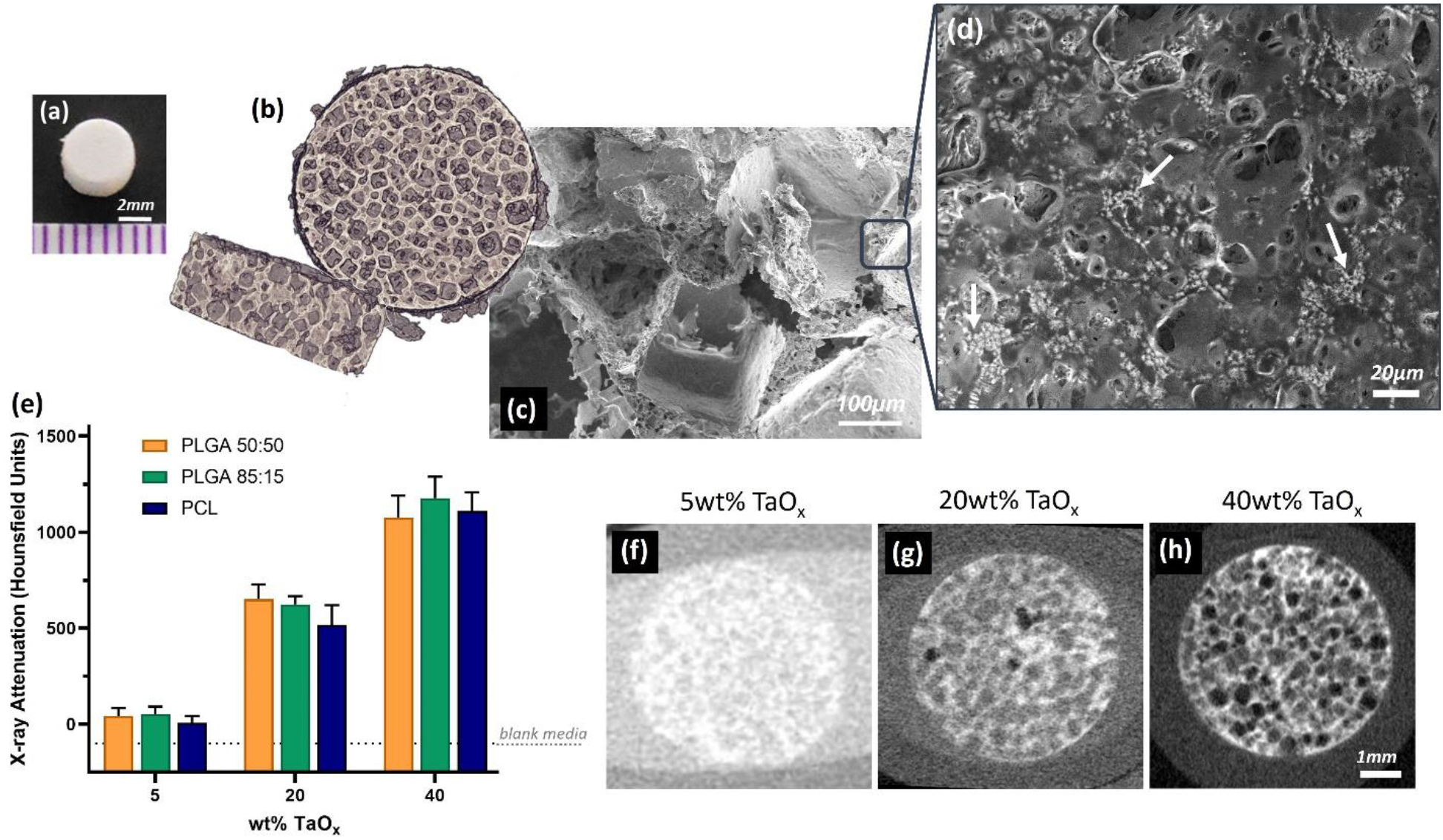
Radiopaque phantoms were produced to mimic key features of implantable biomedical devices. Porous phantoms were disks, as seen (a) macroscopically, containing macro-porous features visible via (b) 3D rendering of a μCT scan and (c) in a corresponding scanning electron micrograph of the phantom cross-section. (d) Radiopaque TaO_x_ nanoparticles were homogeneously dispersed within the polymer matrix (white areas have a high TaO_x_ concentration). (e) X-ray attenuation was dependent on the amount of TaO_x_ incorporated in the matrix, quantified from CT scans of phantoms with (f) 5wt% TaO_x_, (g) 20wt% TaO_x_ and (h) 40wt% TaO_x_. Scale bar (a) 2 mm (c) 100 μm (d) 20 μm (f-h) 1 mm.

### 2.1 Radiopaque Polymer Matrices

Radiopacity was introduced to phantoms via incorporation of hydrophobic TaO_x_ nanoparticles, with a nominal range of 0-40wt% TaO_x_. The addition of nanoparticles to the matrices had no significant effect on the porous phantom structure. A hydrophobic polymer coating on the TaO_x_ particles allowed them to remain suspended and evenly dispersed in the polymer solutions during the manufacturing process. Micrographs of the pore walls show the even dispersion of the TaO_x_(Figure 1(d)), and the homogeneous dispersion ensured that no regions of the polymer matrices had significantly different materials properties or x-ray attenuation. This was critical to ensuring a reproducible degradation profile throughout the structure and that any changes in phantom morphology over time could be observed.

Phantom radiopacity in saline buffer, visualized by micro-computed tomography (μCT), was dependent on the amount of TaO_x_ incorporated into the polymer matrices, in agreement with literature.^[20,21]^ In μCT imaging, radiopacity depends on the ability of a material to attenuate incident x-rays.^[22]^ The higher the concentration of elements, the greater the x-ray attenuation, and the higher the signal. In the case of porous materials like the phantom, the x-ray attenuation measured in the material was a mixture of the radiopaque matrix and the buffer that was trapped within the micro-porous walls. At 5wt% TaO_x_, where phantoms were only just visible (Figure 1(e-f)), the radiopaque matrix could not be distinguished radiographically from the buffer, and the measured radiopacity is a combination of both. Thus, 5wt% TaO_x_ was determined to be the minimum addition that could still allow the gross phantom morphology to be distinguished in a hydrated environment. At higher weight percentages of TaO_x_, sufficient contrast existed between the matrix and the buffer (Figure 1(g-h)), that the micro-porous matrix could be radiographically distinguished from the aqueous media in the macro-pores and segmented by a semi-automated image analysis program. The upper range of TaO_x_ was capped at 40wt% to ensure that the polymer matrix was still the dominant phase within the phantoms. This range of radiopaque nanoparticles is in line with other studies utilizing elements such as gold ^[20]^ and gadolinium.^[21]^The true range of incorporation, after quantification via thermogravimetric analysis (TGA), was found to be 0 - 30 wt% TaO_x_. The difference between theoretical and actual TaO_x_ content is likely due to loss of nanoparticles during the washing step of the leeching process. Regardless, phantoms made with different polymers did not have a significant difference in the amount of TaO_x_ incorporated at each nominal value (± 0.5wt%), which was sufficient to make comparisons between the behavior of the phantoms based on nanoparticle addition.

### 2.2 Degradation profiles

Implantable biomedical devices are designed to support regenerating tissue during the initial stages of healing, and should be engineered to degrade over time, allowing a new functional tissue matrix to replace the implant. Maintaining structural integrity of the device during this time frame is essential, to ensure that infiltrating cells and capillaries are not dislodged or crushed during normal physiological movement of the patient. Defining the amount of time implanted devices must maintain integrity, and thus the corresponding optimal degradation rate for an implanted device, is difficult as it depends on the location of the implant, tissue to be regenerated and overall size of the defect. Compounding the problem is that literature values for in vitro degradation tend to underestimate the in vivo degradation rate, despite being performed under simulated physiological conditions (phosphate buffered saline (PBS), 37°C).^[16]^ These differences are due to the complexity of the in vivo extracellular environment, including blood/lymphatic flow and proteases present within tissue. In addition, the in vivo environment is altered after surgical implantation due to acute inflammation processes, leading to local changes in pH and cell counts. Even when using well characterized, FDA-approved biocompatible polymers, like PCL and PLGA, subtle changes like polymer end groups and chain length can affect the degradation rate and materials properties of the implant during their degradation profile.^[14,15,23]^ The addition of radiopaque nanoparticles to the polymer matrix also have the potential to change the degradation profile, and thus radiopacity of the matrix must be balanced along with other materials properties.

The present study utilized phantoms from three biocompatible polymers that exhibited marked differences in degradation profile: non-degrading (PCL), slow degrading (PLGA 85:15) and fast degrading (PLGA 50:50). This allowed the investigation and tracking of a full range of properties and behaviors and their time course. At one side of the spectrum, PCL with 0wt% TaO_x_ maintained structural integrity for a full 20 weeks, at physiological pH (7.4) and temperature (37°C), where structural integrity is defined as maintenance of a micro-porous structure and > 80% intact gross implant volume. Given the long time scale for PCL hydrolysis, macroscopic changes to PCL scaffolds around physiological pH generally don’t occur prior to 6 months, as corroborated in the current study.^[17]^ PLGA 85:15 also maintained structural integrity at 20 weeks, with 89.6 ± 5.4% of mass remaining. Conversely, the structural integrity of PLGA 50:50 (0wt% TaO_x_) was lost within 11 weeks, corresponding to > 50% mass loss, in line with other reports on isotropic porous devices.^[16]^

To begin to address the issues with extrapolation of in vitro degradation profiles to in vivo environments, phantoms (0-40wt% TaO_x_) were further incubated under pH conditions that simulated chronic inflammation or tumor acidosis (sodium citrate, pH 6.5) ^[13,24]^ and cellular lysosomal conditions (sodium citrate, pH 5.5).^[25]^ As expected, degradation was increased significantly at acidic pH due to increased efficiency of hydrolysis reactions, responsible for the majority of degradation in PLGA and PCL.^[14,17,26]^ The increase in the degradation rate was on the order of weeks, which was not noticeable for PCL within the 20 week period of the study, but significant for the fast-degrading PLGA 50:50, **Figure 2**. As seen visually in Figure 2(a-b), structural integrity at pH 5.5 was lost several weeks before physiological pH. The visible trend correlated with accelerated mass loss in PLGA 50:50 at pH 5.5 compared to pH 6.5 or 7.4, Figure 2(d-f). The effects of buffer pH were maintained regardless of the amount of TaO_x_ included in phantom. Since the radiopaque element did not alter the basic response of the matrices to buffer pH, there is potential to gain critical real time assessment of matrix degradation. This may prove particularly important for “smart” devices that are designed to allow drug release during matrix dissolution, particularly in microenvironments where pH may not be definitively known, such as in tumors where acidosis is prevalent.^[24]^

**Figure 2.**
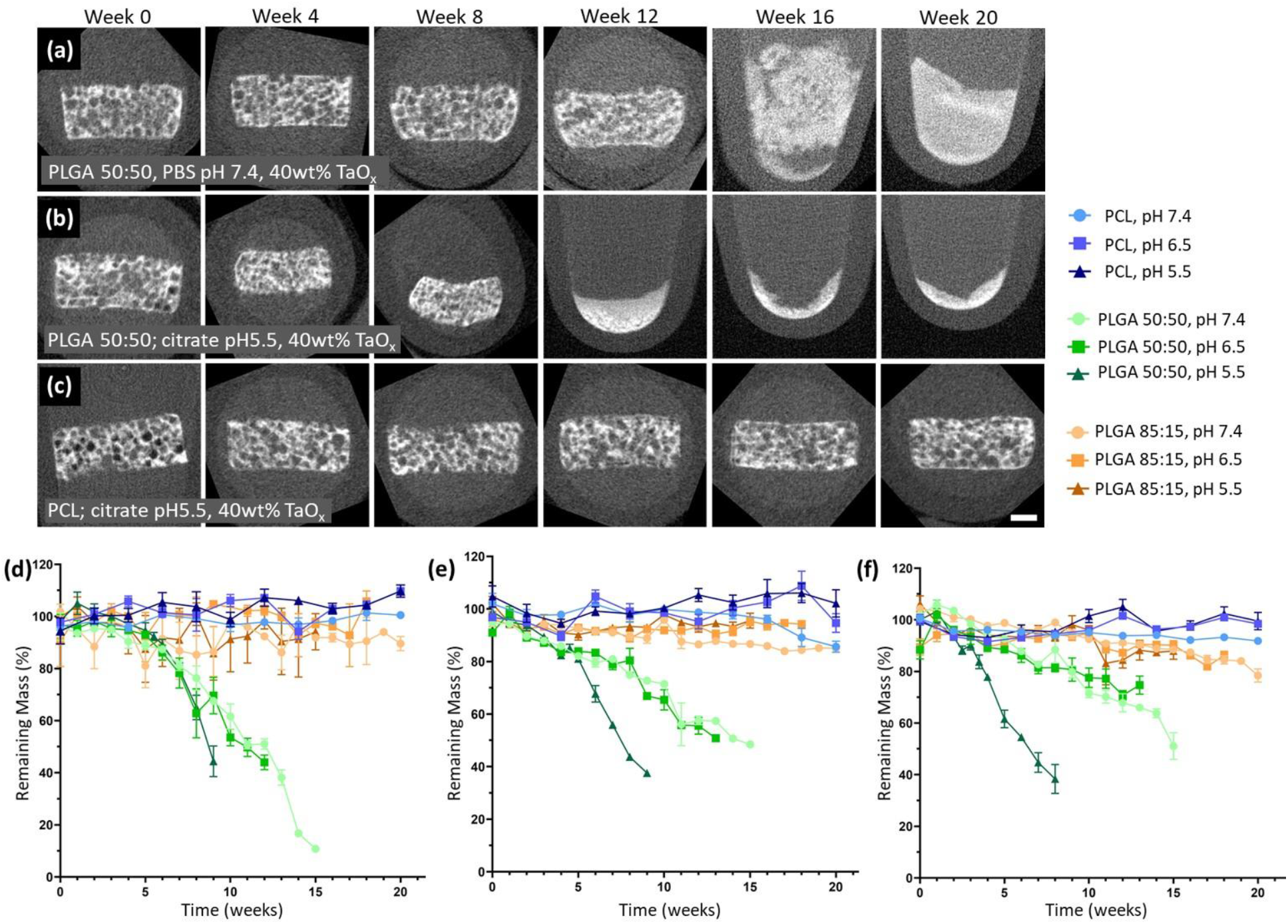
Structural integrity of phantoms over time was largely determined by the matrix polymer, but was lost earlier in lower pH environments. The effects were most noticeable in fast degrading polymers compared to non-degrading polymers, as seen visually in μCT reconstructions over 20 weeks for: (a) PLGA 50:50 + 40wt% TaO_x_ at pH 7.4, (b) PLGA 50:50 + 40wt% TaO_x_ at pH 5.5 and (c) PCL + 40wt% TaO_x_ at pH 5.5. Mass loss mirrored the trends in structural integrity, and was consistent regardless of TaO_x_ incorporation: (d) 0wt% TaO_x_, (e) 20wt% TaO_x_ and (f) 40wt% TaO_x_; data presented as mean ± SEM. Scale bar (a-c): 1 mm.

Not only is structural integrity and matrix mass important to understanding device degradation, but mechanical properties are also key for device function. Both PLGA 50:50 and 85:15 phantoms had significantly higher mechanical strength, measured as a compressive modulus, than PCL initially. Over time, the effects of the buffer pH became far more significant in determining the overall mechanical behavior. In all cases of TaO_x_ incorporation into the phantoms, mechanical stability was lost first at the lowest pH, consistent with the accelerated mass loss. With a high level of porosity within phantoms, the mechanical strength reported is an apparent strength of the structure rather than a true materials property of the matrix. When exposed to a compressive loading, structural collapse of hydrated scaffolds occurs in stages, starting with reversible deformation of the porous walls (giving an apparent compressive modulus), followed by a collapse of the macro-porous structure and a further densification of the remaining micro-pores. As this apparent modulus is dependent on the porosity within the structure, any changes to the apparent density from the degradation process affect the compressive modulus.^[27]^ In particular, PLGA 50:50 experienced the greatest increases in apparent compressive modulus during degradation due to macroscopic changes in overall volume, **Figure 3**, that were influenced by the buffer pH, but PLGA 85:15 also experienced significant increases in strength over the first 10 weeks as well. While PCL had the lowest apparent modulus, mechanical strength was maintained over the 20 week period.

**Figure 3.**
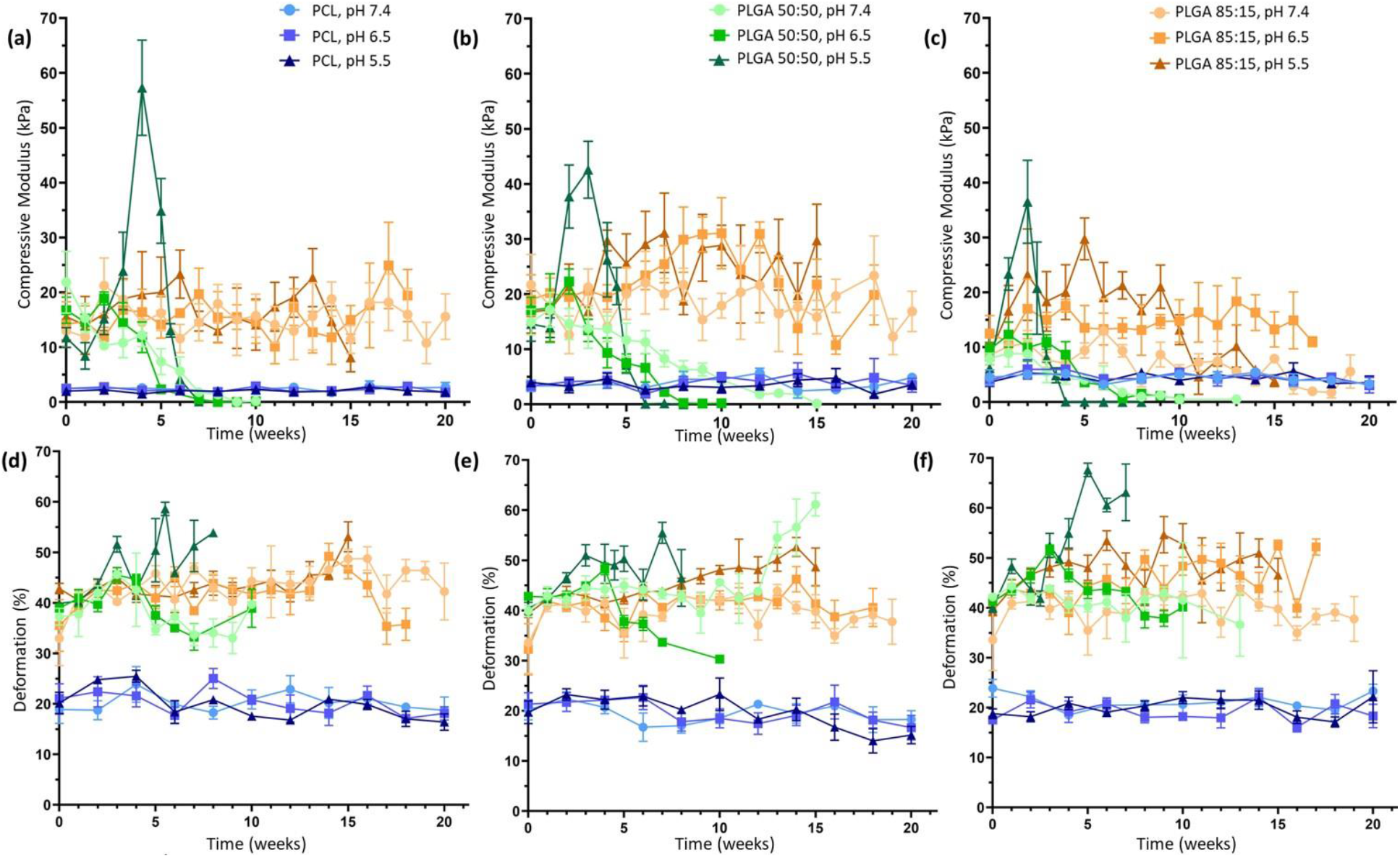
The polymer matrix in the porous phantoms drove the mechanical behavior, as measured by (a-c) apparent compressive modulus and (d-f) deformation. The pH of the buffer had a significant impact on mechanical properties at all levels of TaO_x_ incorporation: (a, d) 0wt% TaO_x_ (b, e) 20wt% TaO_x_ and (c, f) 40wt% TaO_x_. Data reported as mean ± SEM.

Deformation is another feature of devices affecting overall functionality, particularly in cases when devices are implanted in high stress environments, where large deformations after mechanical loading correlate to a permanent loss of a devices pre-stressed shape and potential catastrophic failure. The deformation of the phantoms, measured as the recovery of phantom thickness after compression ^[28]^, was dependent on both the polymer matrix and the pH of the buffer. Deformation of PCL phantoms was consistently around 20%, demonstrating a high level of compliance and recovery. In contrast, PLGA phantoms behaved as brittle materials, with very little recovery in the micro-porous structure.^[28]^ In general, deformation tended to increase as densification of the scaffold occurred, particularly at low pH, and changed significantly as the mechanical strength was lost, **Figure 4**(d-f).

**Figure 4.**
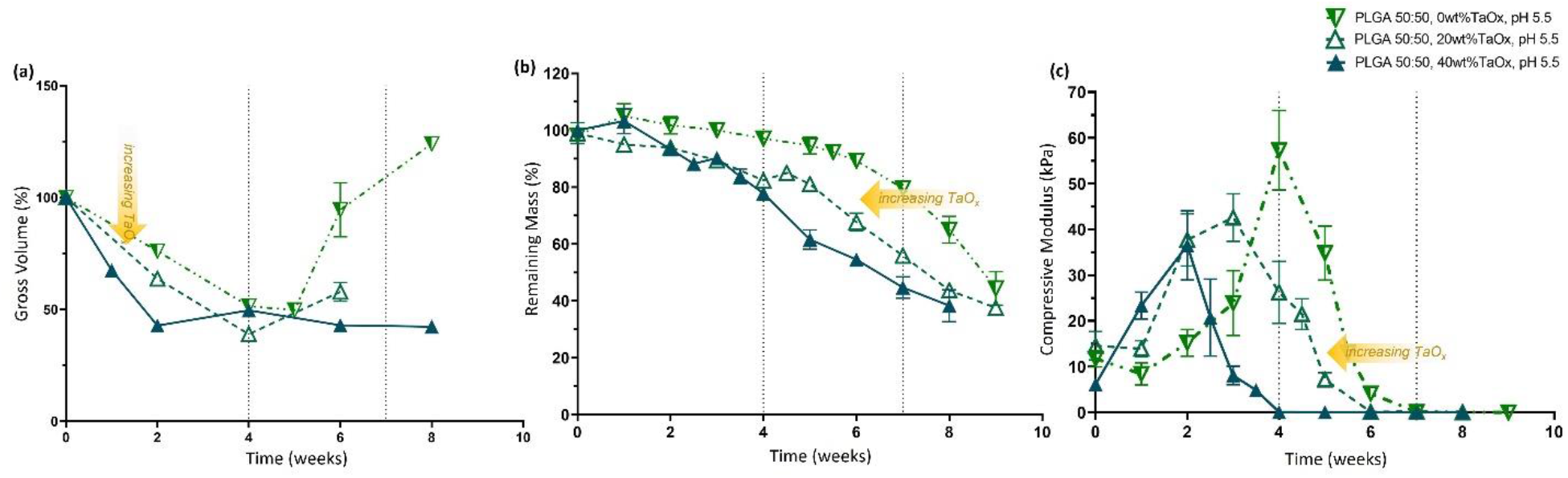
TaO_x_ incorporation increased phantom hydrolysis in fast-degrading PLGA 50:50. At pH 5.5, increased wt% TaO_x_ sped up the rate of (a) initial volume change, (b) mass loss and (c) loss of compressive modulus over time. Metrics for quantifying degradation do not share the same features. Data reported as mean ± SEM.

Importantly for understanding degradation profiles of biomedical devices, different metrics responded at significantly different time scales. Phantoms from PLGA 50:50 demonstrated consistent loss of mechanical stability without a correspondingly significant change in mass loss, Figure 4. Both mass loss and mechanics are often reported as measures of matrix stability during degradation, but there is little predictive value between them. Indeed, the processes governing the two metrics are different. Mechanical strength is affected by cleavage of polymer chains within the porous structure, and in this case, significant changes to apparent density. However, mass loss has been compared to an erosion process, which is dependent on both cleavage of polymer chains within the structure and the diffusion of material from the implant itself.^[14]^ Thus, the solubility of degradation products, swelling and surface area can all affect mass loss in ways that may not be as important for mechanical integrity.^[14]^

While trends in the buffer pH on degradation remained consistent regardless of nanoparticle incorporation, the addition of nanoparticle fillers into polymer matrices is also known to independently affect the mechanical strength and degradation of composite matrices.^[29]^ In the case of the porous phantoms, there were trends in the compressive modulus of the matrices, as the TaO_x_ content increased to 40wt%. In a PCL matrix, the nanoparticles tended to create a stiffer matrix, likely hindering the sliding of polymer chains within the pore walls; apparent modulus of the phantoms went from 2.59±1.2 to 4.51±2.7 kPa with 0 and 40wt% respectively. In contrast, both types of PLGA matrices had reduced mean compressive modulus as the incorporation of TaO_x_ increased from 20wt%; the apparent modulus decreased by up to half for PLGA 50:50 (15.91 ± 11.05 to 7.90 ± 1.99 kPa). Interactions between the polymer chains and the hydrophobic layer around the nanoparticles may not allow entanglement of the individual polymer chains, thus facilitating movement of the pore walls. Despite the trends, the effect of the porosity on the phantom modulus, both micro and macro, created too much variation for statistical significance.

Effects from TaO_x_ incorporation over time appeared to be mediated by increased water penetration into the polymer matrix of the phantoms, increasing hydrolysis reactions. In the case of PLGA 50:50, all metrics describing scaffold integrity and mechanical stability experienced faster decline with increased weight percent of TaO_x_, Figure 4. Loss of mechanical strength for PLGA 50:50 at pH 5.5 occurred after 4 weeks for 40wt% TaO_x_ as opposed to 7 weeks for native polymer (0wt% TaO_x_), Figure 4(c). In PLGA 85:15, an increased hydrolysis correlated to an increase in the rate of volume changes within the scaffold matrix, if not a clear correlation with loss of mechanics. PCL phantoms, in contrast, appeared largely unaffected by the addition of nanoparticles. In other systems, PCL composite materials have been shown to have increased degradation rates, due to the added intercalation of water within the matrix, effectively increasing the surface area available for hydrolysis.^[29]^ However, in these studies the size of the particles and porosity of the matrix was completely different, making direct comparison difficult.

From the present study, it is clear that degradation is largely controlled by the polymer matrix used to create the phantom. Local environmental factors, such as pH, played a much larger role in accelerating degradation processes than the addition of radiopaque nanoparticles. Considering the traumatic environment encountered by biomedical devices upon implantation, further acceleration of degradation with large amounts of TaO_x_ (40wt%) would likely be impractical for most applications, creating an effective upper limit on TaO_x_ incorporation. The TaO_x_ nanoparticles would likely shorten PCL degradation time as well, but whether on the order of weeks or months is currently unclear.

### 2.3 Longitudinal Monitoring

To translate longitudinal monitoring to the clinic successfully requires the ability to clearly track changes in biomedical devices over time, despite damage or disease in surrounding tissues. Thus, any potential monitoring paradigms must be robust during both mechanical failure, device movement and degradation, which represent common failure mechanisms of biomedical devices in diverse areas from hernia patches to knee replacement.^[1,30,31]^ With the promising results on phantom radiopacity at initial time points, we further demonstrated that the incorporation of TaO_x_ nanoparticles resulted in radiopaque phantoms that could be imaged through a 20 week time course, Figure 2. Importantly, the TaO_x_ appeared to remain associated with the polymer matrix, despite loss of structural integrity of the phantoms, allowing for real time characterization of the phantoms which could corroborate the data on mechanical properties and mass loss.

Since the x-ray attenuation over time was dependent on the amount of TaO_x_ present in the matrix, only the gross volume, or the volume occupied by a solid cylinder with the corresponding outside thickness and diameter measured by CT, could be recorded for phantoms with 5wt% TaO_x_, **Figure 5** (d, e1, f1). Even from the gross volume alone, very clear trends in the phantom volume changes over the degradation period were visible. The changes in volume were driven primarily by the buffer media and from degradation products such as lactic acid, that could affect the environment within the phantom itself.^[14]^ It is well known that PLGA will experience bulk degradation, caused by the breakdown of the glycolic acid chain and release of lactic acid that has auto-catalytic activity by lowering the local pH within porous structures. As expected, with its high glycolic acid content, PLGA 50:50 was far more affected during degradation than PLGA 85:15, evidenced by drastic changes in gross volume.^[16]^ At a buffer pH of 5.5, PLGA 50:50 rapidly contracted to < 50% its original gross volume, consistent with other studies that note volume decreases coincident with pH drops within devices during degradation.^[27]^ In this case, the buffer was responsible for the acid environment rather than the acid by-products of degradation. Conversely, at a buffer pH of ≥ 6.5, PLGA 50:50 phantoms tended to swell. It has been hypothesized that the internal build up of acidic by-products encourages water infiltration into PLGA structures and relaxation of the polymer matrix which further drives the swelling behavior.^[32]^ Peak swelling occurred at pH 6.5, reaching >250%, >600% and >150% of original scaffold volume for 5, 20 and 40wt% respectively. These extreme values for swelling are likely underestimated for 5wt%, as analysis was conducted on CT scans, and 5wt% could not be distinguished from the background after 6 weeks, even for gross volume calculations. CT scans of the radiopaque PLGA 50:50 noted the development of cavitation in the interior of the phantom matrix over time at pH 6.5 and 7.4, while the outer surface remained intact. This is typical of internal bulk degradation of PLGA constructs, where there is little outward indication of the loss of structural integrity from macroscopic observation.^[14]^ Like PLGA 50:50, PLGA 85:15 phantoms also underwent significant volume changes, namely contraction, over the study period, which was most evident at pH 5.5. A typical time frame for PLGA 85:15 devices to exhibit gross morphology changes is around 19-24 weeks, in line with the current study.^[16,27]^

**Figure 5.**
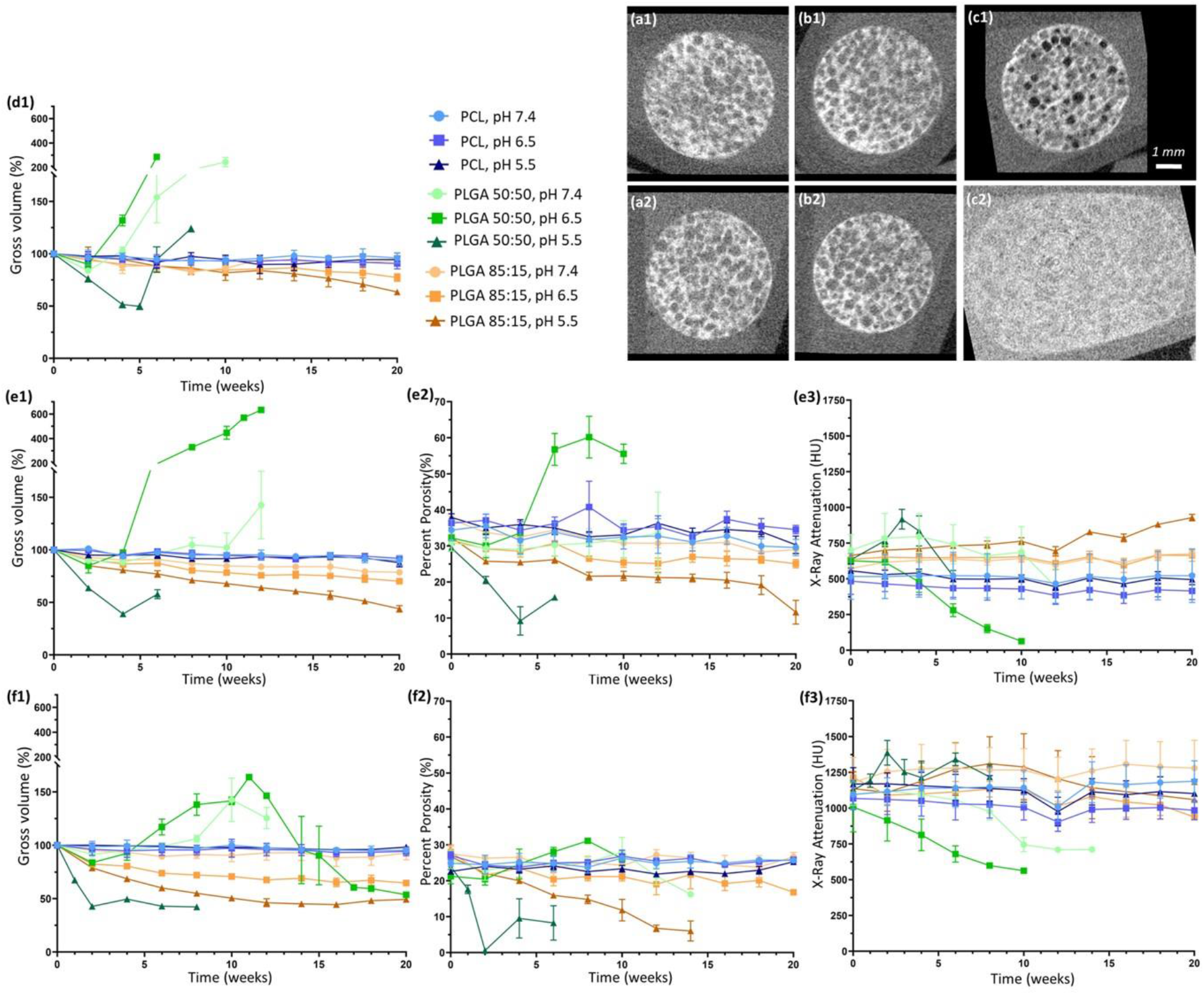
Phantoms with TaO_x_ nanoparticles could be monitored for 20 weeks without loss of radiopacity due to particle leeching. (a-c) This allowed for visual monitoring of changes to phantom shape and porosity, as illustrated by 3D reconstructions from (1) day 1 and (2) 6 weeks: (a) PCL+20wt% TaO_x_ (b) PLGA 85:15 + 20wt% TaO_x_ and (c) PLGA 50:50 + 20wt% TaO_x_. During degradation, significant changes occurred within the phantoms: (1) gross phantom volume (2) percentage porosity and (3) x-ray attenuation. TaO_x_ incorporation ranged from (d) 5wt% TaO_x_, (e) 20wt% TaO_x_ and (f) 40wt% TaO_x_. At 5wt% TaO_x_, only gross volume of the phantoms could be quantified, as the matrix could not be segmented from the background. Scale (a-c): 1 mm.

The effect of degradation on the porous structure could be examined more closely in phantoms with >5wt% TaO_x_, as the volume of the matrix could be separated from the internal macro-pores. Percent porosity, related to the macro-pores within the phantom, was calculated as the percentage of the gross volume not occupied by matrix. Prior to degradation, all scaffolds with 20wt% TaO_x_ had between 30-40% porosity, and all 40wt% TaO_x_ phantoms had a 20-30% porosity, Figure 5(e2, f2). The difference between the two groups in initial porosity is an overestimation of the matrix volume during the segmentation process, the result of balancing the threshold for the program to distinguish matrix from background consistently through the entire series of phantoms. From the percentage porosity, it was clear that swelling of PLGA 50:50 at pH 6.5 increased the internal porosity significantly, while the densification of phantoms at pH 5.5 appeared to be driven by a decrease in open pore volume over and above decreases in polymer matrix volume. This implies a restriction of nutrient or fluid flow through the porous phantoms as degradation progresses, or even a total collapse of porosity in the worst case scenario. The extreme contraction of porous devices during cellular infiltration has been tied to poor repair outcomes, necessitating research into ways to resist contractile forces.^[33,34]^ Swelling, on the other hand, is not likely to interfere with tissue repair, as biomedical implants are generally constrained by surrounding tissues after implantation. Swelling might, however, influence diffusion of drugs or therapeutics from a polymer matrix.^[32]^

As observed in the initial characterization of radiopaque phantoms, the x-ray attenuation was dependent on the amount of TaO_x_ present in the matrix. While the matrix remained at a constant volume, as was the case for PCL throughout the study, the x-ray attenuation remained unchanged. On the other hand, as PLGA matrices contracted or expanded, the x-ray attenuation of the phantoms inversely increased and decreased, Figure 5(e3, f3). This is largely due to the apparent concentration of the contrast agent, TaO_x_ in this case, within the matrix. As there was no significant change in TaO_x_ amount, movement of the matrix either concentrated or diluted the nanoparticles which were available to attenuate x-rays. At pH 6.5, PLGA 50:50 with 5 or 20wt% TaO_x_ expanded to such an extent (Figure 5(ed1, e1)) that there was no longer sufficient contrast to either visualize the scaffold or segment the matrix from the buffer, due to the drop in attenuation. In contrast, at pH 5.5, PLGA 50:50 phantoms showed increases in phantom attenuation before the structural integrity was lost, Figure 5(e3, f3). Even after integrity was lost, enough TaO_x_ remained associated with the polymer to distinguish the remains of the phantom structure from the surrounding environment, Figure 2(b).

As in the case of phantom mechanics, nanoparticle incorporation had an independent effect of the buffer pH on volume changes. Overall, higher percentages of TaO_x_ tended to lower the overall volume of the phantom. This manifested as either an increasing phantom contraction or decreased swelling. In PLGA phantoms, this also manifested as an increase in the rate of volume change, most apparent for PLGA 85:15. With the known bulk degradation mechanism of PLGA, the higher levels of TaO_x_ may limit the build up of acidic degradation products within the scaffold core, which has shown to be tied to swelling kinetics in PLGA microspheres.^[32,35]^ The nanoparticles may be contributing by limiting the amount of degradation product available overall or by increasing the hydrolysis of the polymer matrix to allow the escape of acidic groups before significant build up.

Within the in vitro microenvironment, the nanoparticles remain associated with the matrix rather than remaining suspended in the degradation buffer, likely due to the hydrophobic nature of the polymer and nanoparticles. During in vivo degradation, both polymer degradation products (monomers, oligomers) and TaO_x_ nanoparticles are expected to behave in a similar manner. They must diffuse to the blood, after which they can be excreted via liver, kidneys and spleen.^[9,36]^ For nanoparticles, clearance has been shown to go rapidly from the blood, with less than 1.15% injected dose remaining in the body after 48 hrs.^[9]^ Previous in vivo nanoparticle studies have not demonstrated any blurring of device features during sequential CT scans due to dispersed nanoparticles, so the results in long-term monitoring should be consistent with the data collected during in vivo applications.^[21]^

The incorporation of nanoparticles within slow degrading polymer matrices is a way to conveniently localize radiopaque markers or implants, bypassing the immediate problems with cellular toxicity of concentrated nanoparticle bolus injections and providing a sustained signal.^[11,20,21]^ Introducing radiopacity to implantable devices is a facile and flexible way of monitoring and predicting implant failure, particularly where local tissue environments are in flux due to trauma or disease.^[3]^ A range of 5-20wt% TaO_x_ appears optimal for synthetic biodegradable scaffolds, enabling visualization without significantly impairing mechanical properties. Due to differences in inflammatory responses and mechanical forces at different implantation sites, the physiological environment for every implant is expected to be slightly different. Thus, having information on the full range of potential degradation profiles is critical for the rational design of next generation implants. Differential location within the body is not a concern for CT imaging, as penetration of high energy x-rays have excellent resolution with no limitations on penetration depth.^[2]^ Indeed, phantoms with 20-30 wt% TaO_x_ could be implanted and successfully monitored when placed subcutaneously, intraperitoneally and intramuscularly in a rodent model, **Figure 6**, and movement over time of the implants could be easily tracked. This clearly offers the flexibility needed for biomedical implants, while working within already established clinical workflows for imaging patients.

**Figure 6.**
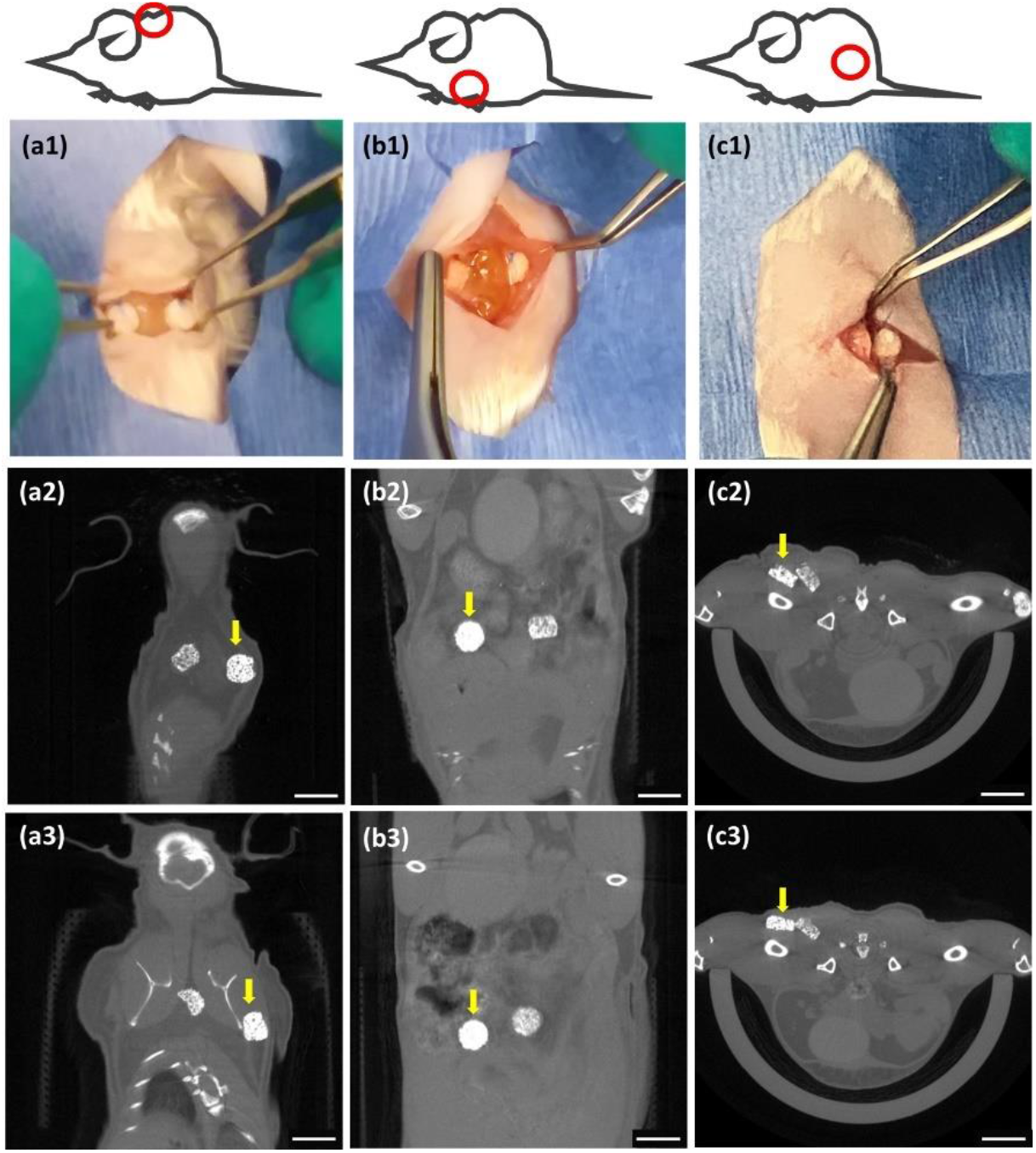
In situ monitoring using CT is possible in a variety of locations including (a) subcutaneous, (b) peritoneal and (c) intramuscular. Implantation of phantoms containing 20 and 30wt% TaO_x_ as viewed (1) macroscopically at the time of implantation into mice and via μCT scanning at (2) day 1 post-implantation and (3) 5 weeks post-implantation. Movement of the scaffolds is noted over time, particularly in the subcutaneous space (a1-3). Each implanted phantom was 3 mm diameter x 1.5mm thick; yellow arrow marks the scaffold incorporating 30wt% TaO_x_. Scale bar (2-3): 5 mm.

## 3. Conclusions

Radiopaque implantable devices offer the potential to monitor the real time response of polymeric devices to their environment and to predict device failure, both during device development phase and in the clinic. Hydrophobic radiopaque TaO_x_ nanoparticles were incorporated homogeneously within a variety of biocompatible polymer phantoms, with 5wt% TaO_x_ being the minimum to enable in situ monitoring of gross phantom features (overall volume, location) using μCT. Beyond 20wt% TaO_x_, there was limited advantage in terms of imaging, and increasing destabilization of the polymer matrix. Importantly, within this range of 5-20wt%, radiopacity of phantoms was maintained over 20 weeks. Phantom x-ray attenuation was impacted most significantly by swelling and contraction during degradation, which essentially diluted or concentrated the contrast agent. The overall degradation profile of phantoms was dictated by the polymer matrix. Lower pH environments and high nanoparticle content (> 20wt% TaO_x_) increased hydrolysis, simultaneously increasing degradation rate, as measured by mass loss, structural integrity and mechanical stability. This study demonstrates a comprehensive testing of phantom degradation profiles with simultaneous imaging, representing a significant step towards incorporating in situ monitoring into the next generation of implantable devices.

## 4. Materials & Methods

The study utilized three types of biocompatible polymers: polycaprolactone (PCL), poly(lactide-co-glycolide) (PLGA) 50:50 and PLGA 85:15. PCL (Sigma Aldrich) had a molecular weight average of 80 kDa. PLGA 50:50 (Lactel/Evonik B6010-4) and PLGA 85:15 (Expansorb^®^ DLG 85-7E, Merck) were both ester terminated and had a weight average molecular weight of between 80-90 kDa, to minimize the effects of polymer chain length on the degradation rate.^[14]^

### 4.1 Hydrophobic TaO_x_ nanoparticles

Hydrophobic TaO_x_ nanoparticles were manufactured based on our previous procedure with one minor change.^[11]^ Hydrophobicity is imparted by coating nanoparticles with aliphatic organisilanes. Here, the organosilane used was hexadecyltriethoxysilane (HDTES, Gelest Inc, cat no SIH5922.0) rather than (3-aminopropyl) trimethoxy silane (APTMS, Sigma).

### 4.2 Phantom manufacture

Phantoms were created with micro-scale porosity (< 100μm) and with macro-scale porosity (200 - 500 μm) to mimic tissue engineering constructs which must accommodate both nutrient diffusion and cell and tissue infiltration. Biocompatible polymers were solubilized in suspensions of TaO_x_ nanoparticles in dichloromethane (DCM, Sigma); solutions were 8wt% and 12wt% for PCL and PLGA, respectively. TaO_x_ suspensions were calculated so that the final phantom mass (polymer + nanoparticles) would consist of 0 - 40wt% TaO_x_. Sucrose (Meijer) was added to the suspension, calculated to be 70 vol% of the polymer + nanoparticle mass in solution, followed by NaCl (Jade Scientific) at 60 vol% of the total polymer + nanoparticle volume. The suspension was vortexed for 10 minutes and pressed into a silicon mold that was 4.7mm diameter, 2 mm high. After air drying, phantoms were removed, trimmed of excess polymer and then washed for 2 hours in distilled water, changing the water every 30 minutes to remove sucrose and NaCl. Washed phantoms were air dried overnight and stored in a desiccator prior to use.

### 4.3 Scanning electron microscopy

Samples were adhered to 13mm aluminum stubs and sputter coated with platinum. Surfaces were examined using a Zeiss Auriga, at 7 keV in scanning electron mode, and 20 keV for backscatter electron images.

### 4.4 Thermogravimetric analysis (TGA)

TGA was used to characterize the weight percentage of TaO_x_ incorporated into polymeric phantoms. Using a TA Q500 (TA Instruments), 12-14 mg of dry material was massed in an alumina pan. The sample was then heated to 600°C at a rate of 10°C/min under a nitrogen environment. The percentage of nanoparticles was calculated as the difference between the initial and final mass.

### 4.5 Degradation study

An in vitro degradation study was performed to assess the effects of the TaO_x_ incorporation on device properties. Three buffers were used, to simulate a range of physiological conditions: pH 7.4 phosphate buffered saline (PBS, Thermo Fisher Sci), pH 6.5 0.05 M sodium citrate (Thermo Fisher Sci), and pH 5.5 0.05 M sodium citrate (Thermo Fisher Sci). Dry phantoms were placed into individual, pre-weighed, microcentrifuge tubes. Phantoms were hydrated on day 0, by adding 1 ml buffer and centrifuging for 10 min at 11,000 rpm. Afterwards immersed phantoms were stored at 37°C, and the buffer was changed weekly. Phantoms with 0, 20 and 40wt% TaO_x_ were tested for mechanics and mass loss (n=4 per time point). A further cohort of phantoms (n=3) containing 5, 20 and 40wt% TaO_x_ were serially imaged via μCT, to follow the degradation process and assess radiopacity over time.

### 4.6 Mechanics and mass loss

Prior to performing mechanical tests, phantom thickness and diameter was measured using a micrometer. Phantoms were loaded onto a TAXT plus Texture analyzer (Stable Micro Systems) with a 5 kg load cell, compressed to 40% of their original thickness, at 1 mm/min and then unloaded. The loading cycle was repeated twice. The compressive modulus was calculated as the initial slope of the stress-strain curve from the first loading cycle. The percentage of deformation was obtained by dividing the phantom thickness after the initial loading cycle by the original thickness x 100%. The phantom thickness post-loading was determined to be the point at which the force applied to the phantom was greater than 0.02 N during the second loading cycle. After testing, phantoms were removed from the machine, placed in the original microcentrifuge tube and washed in 1 ml deionized water, shaking at 80rpm, for 1 hour at room temperature. The water was subsequently removed and the phantoms were dried at room temperature before determining remaining mass. Mechanics and mass loss for PLGA phantoms were measured at least once per week up to 20 weeks. Phantoms made from PCL, a non-degrading matrix, were measured once every two weeks for 20 weeks.

To calculate mass loss, initial phantom mass was computed as the difference between the empty microcentrifuge tubes and the microcentrifuge tube + phantom. After mechanical testing, dried phantoms were weighed again and the difference from the empty microcentrifuge weight was taken as the remaining mass.

### 4.7 Micro Computed Tomography (μCT)

All tomography images were obtained using a Perkin-Elmer Quantum GX. At every time point, groups of three phantoms were imaged at 90 keV, 88 μA, with a 25 mm field of view at a 50 μm resolution. After acquisition, individual phantoms were sub-reconstructed using the Quantum GX software to 12-18μm resolution. Phantoms used for serial monitoring were imaged on day 0 prior to hydration, and imaged again 24 hrs after hydration with buffer. Through the remainder of the study, all groups were imaged every week after changing the buffer media.

In vivo μCT on mice, was performed at 90 keV, 88 μA, using a 72mm field of view (14 minutes total scan time), 90μm resolution. During acquisition, mice were anesthetized using inhalant anesthetic 1-3% Isoflurane in 1.0L/min Oxygen. Mice were scanned immediately post implantation, on day 1 post-implantation and at day 7 and week 5 post implantation. Total cumulative radiation dosage was less than 24 Gy over 5 weeks.

### 4.8 Tomography image analysis

From the tomography scans of phantoms, several parameters were quantified. Gross features of the phantoms (thickness, diameter) were analyzed using Image J. For phantoms with 5wt% TaO_x_, 50μm scans were used to collect data; 12-18 μm sub-reconstructions were used for all other phantoms. Image stacks were opened and rotated, using built in functions, to measure thickness and diameter in 5 planes, which were averaged. From the diameter and thickness, a “gross volume” was defined as the volume occupied by a solid cylinder with the corresponding thickness and diameter. In the case of the phantoms, this gross volume consisted of both the polymer volume and the volume of the pores.

Analysis of the polymer matrix component of phantoms with 20wt% and 40wt% TaO_x_ was performed using custom software developed with MATLAB (v R2021b, Mathworks, Natick, MA) on μCT sub-reconstructions; phantoms with 5wt% TaO_x_ could not be radiographically distinguished from the background. Before segmenting the polymer from the background, the image was preprocessed by using an adaptive histogram equalization technique to enhance contrast.^[37]^ After, Otsu’s binary segmentation method was used to create a rough mask of the volume.^[38]^ An adaptive thresholding method was then employed to segment the polymer within the rough mask from the background.^[39]^The resulting volume was cleaned up using erosion and dilation operations. The program calculated mean attenuation of the phantom, the volume of the polymer in the phantom, phantom diameter and thickness, number of pores, and average pore diameter. From this, percent porosity of the phantoms was calculated as the percentage of the gross volume not occupied by matrix.

### 4.9 Pilot implantation

PCL implants containing 20wt% and 30wt% TaO_x_ nanoparticles were prepared as detailed above After preparation, phantoms were cut to 3mm diameter with a biopsy punch and were soaked in 70% ethanol for 2 hours. The ethanol was replaced by sterile PBS and sonicated for 5 minutes, twice, and the implants were left in sterile PBS at 37°C until implantation.

All procedures were performed in accordance with IACUC approved protocols and Veterinary guidelines at Michigan State University. BALB/c Mice (N=3 adult male, 7 months old; Charles River Laboratories) were used for this surgical implantation and μCT imaging pilot study. Each mouse was surgically implanted with two PCL implants containing either 20wt% or 30wt% TaO_x_ nanoparticles installed adjacent to each other within the single implantation surgical site. The PCL implants were implanted subcutaneously (N=1 mouse) dorsal-thoracic between the scapulae, intramuscularly (N=1 mouse) dorsal-medially between the biceps femoris and superficial gluteal muscles, and intraperitoneally (N=1 mouse) via a ventral midline laparotomy superficial to the liver. Each animal was administered analgesia at least 20 minutes prior to making the initial incision, including prophylactic Ampicillin (25mg/kg; SID) administered S.C., Meloxicam (5mg/kg; SID) administered S.C. in the right dorsal lateral flank of the animal, and local infiltration of 2% Lidocaine (diluted to 0.5%) was administered S.C. along the intended incision site just prior to making the cutaneous incision (7mg/kg Max dose; SID). Animals were anesthetized via inhalant isoflurane (3-4% isoflurane in 0.8-1LPM oxygen for induction) and maintained via inhalant isoflurane during surgery (1-3% isoflurane in 0.8-1LPM oxygen). Supplemental heat was provided via recirculating warm water blankets during anesthesia induction, patient preparation, surgery, and patient recovery. Each scaffold was sutured to surrounding tissue with at least one single interrupted suture using 9-0 PROLENE^®^ (Polypropylene) monofilament Suture for in-situ location retention. Incision sites were closed in multiple layers where necessary using 5-0 COATED VICRYL^®^ (polyglactin 910) Suture and cutaneous layers closed using 5-0 PDS-II^®^ (polydioxanone) Suture. Following animal recovery, Meloxicam (5mg/kg; SID) and Buprenorphine (2.0mg/kg, BID, every 8-12 hrs) was administered S.C. in the left or right dorsal lateral flank of the animal for 48-hours following surgery. Post-Operative clinical observation, body weight assessment, and health score assessment were performed for 7-14 days post-operatively. Animals were scanned with μCT immediately post implantation, and again on day 1, day 7, and week 5 post implantation. At termination, animals were sacrificed via CO2 overdose asphyxiation.

### 4.10 Statistics

Statistics were performed using GraphPad 9.4.1. Mechanical data was analyzed via non-parametric Kruskal-Wallis test followed by multiple comparisons of the means, using two-stage step-up method of Benjamini, Krieger and Yekuteli test, to control the False Discovery Rate. All other data was analyzed via ANOVA, followed by Tukey’s multiple comparisons test. In all cases, α < 0.05 was considered significant, with a 95% confidence interval.

## Acknowledgements

The authors would like to thank P. Askeland from the MSU Composite Materials and Structures Center for help with scanning microscopy. This study was funded by the National Institute of Biomedical Imaging and Bioengineering of the NIH under award number R01EB029418. The content is solely the responsibility of the authors and does not necessarily represent the official views of the National Institutes of Health.

## Notes

### Competing Interest Statement

The authors have declared no competing interest.

